# AANAT1 functions in astrocytes to regulate sleep homeostasis

**DOI:** 10.1101/736223

**Authors:** Sejal Davla, Gregory Artiushin, Daryan Chitsaz, Sally Li, Amita Sehgal, Donald J. van Meyel

## Abstract

Characteristic features of sleep are conserved among species [1], and from humans to insects sleep is influenced by neural circuits involving monoamines such as serotonin and dopamine [2]. Glial cells have been increasingly implicated in mechanisms of baseline and homeostatic sleep regulation in mammals and flies [3–11], but it remains unknown whether and how glia might influence monoaminergic control of sleep. Sleep is regulated by circadian rhythms and a homeostatic drive to compensate for prolonged wakefulness, and growing evidence suggests that neural mechanisms controlling homeostatic sleep can be discriminated from those controlling baseline sleep [12–15]. In *Drosophila*, mutants of arylalkylamine N-acetyltransferase 1 (*AANAT1^lo^*) have normal baseline amounts of sleep and motor activity, but increased rebound sleep following deprivation [16]. AANAT1 can acetylate and inactivate monoamines *in vitro* [17], but the role of AANAT1 *in vivo* remains poorly understood. We find AANAT1 to be expressed in astrocytes and subsets of neurons in the adult *Drosophila* brain, with levels in astrocytes declining markedly overnight. In sleep-deprived *AANAT1* mutant flies, heightened rebound sleep is accompanied by increased serotonin and dopamine levels in the brain. In neurons, AANAT1 functions to limit the quantity and consolidation of nighttime sleep, but in astrocytes AANAT1 constrains the amount of rebound sleep that flies take in response to sleep deprivation. These findings distinguish sleep-control functions of AANAT1 in neurons and astrocytes, and identify a critical role for astrocytes in the regulation of monoamine bioavailability and calibration of the response to sleep need.

**Highlights:**

- The monoamine catabolic enzyme arylalkylamine N-acetyltransferase 1 (AANAT1) is expressed by astrocytes and subsets of serotonergic, glutamatergic, GABAergic and cholinergic neurons in the adult brain of *Drosophila*.
- AANAT1 limits accumulation of serotonin and dopamine in the brain upon sleep deprivation.
- Loss of AANAT1 from astrocytes, but not from neurons, causes flies to increase their daytime rebound sleep in response to overnight sleep deprivation.

## Results and Discussion

We generated antiserum to AANAT1 (known previously as Dopamine acetyltransferase (Dat) and confirmed its specificity with immunohistochemistry (IHC) in the embryonic CNS. AANAT1 immunoreactivity was observed in the cytoplasm of many cells (**Fig. 1A**) but was absent in age-matched embryos that were homozygous for a deletion of the entire AANAT1 gene (**Fig. 1B**). In adult brains, AANAT1 was co-labeled with Bruchpilot (nc82, **Fig. 1C**), a presynaptic marker that labels neuropil regions. We found that AANAT1 was expressed in neurons (Elav^+^, **Fig. 1D,D’**) and glia (Repo^+^, **Fig. 1E,E’**) throughout the brain. With Gal4 drivers specific for glial subtypes, we confirmed that AANAT1-positive (^+^) glial cells are astrocytes. All astrocytes (Alrm-Gal4) labeled by a Red Fluorescent Protein (RFP) reporter (Alrm>nuRFP) were AANAT1^+^ in the central brain (**Fig. 1F-F’’**). With pan-neuronal RNAi-mediated knockdown of AANAT1 using the driver nSyb-Gal4 and two distinct RNAi lines, the AANAT1-positive glial cells could be identified more clearly as astrocytes by their ramified morphology and the presence of AANAT1 in their thin processes that infiltrate neuropil (**Fig. S1**). Finally, their identity was further confirmed by the morphology of single cells with Multi-Color Flp-OUT (MCFO) analysis (**Fig. 1G**). Only a subset of astrocytes in the optic lobes residing between the medulla and lobula did not express AANAT1 (**Fig. S1**). AANAT1 expression was absent from ensheathing glia marked by R56F03-Gal4 (**Fig. S1**).

**Figure 1.**
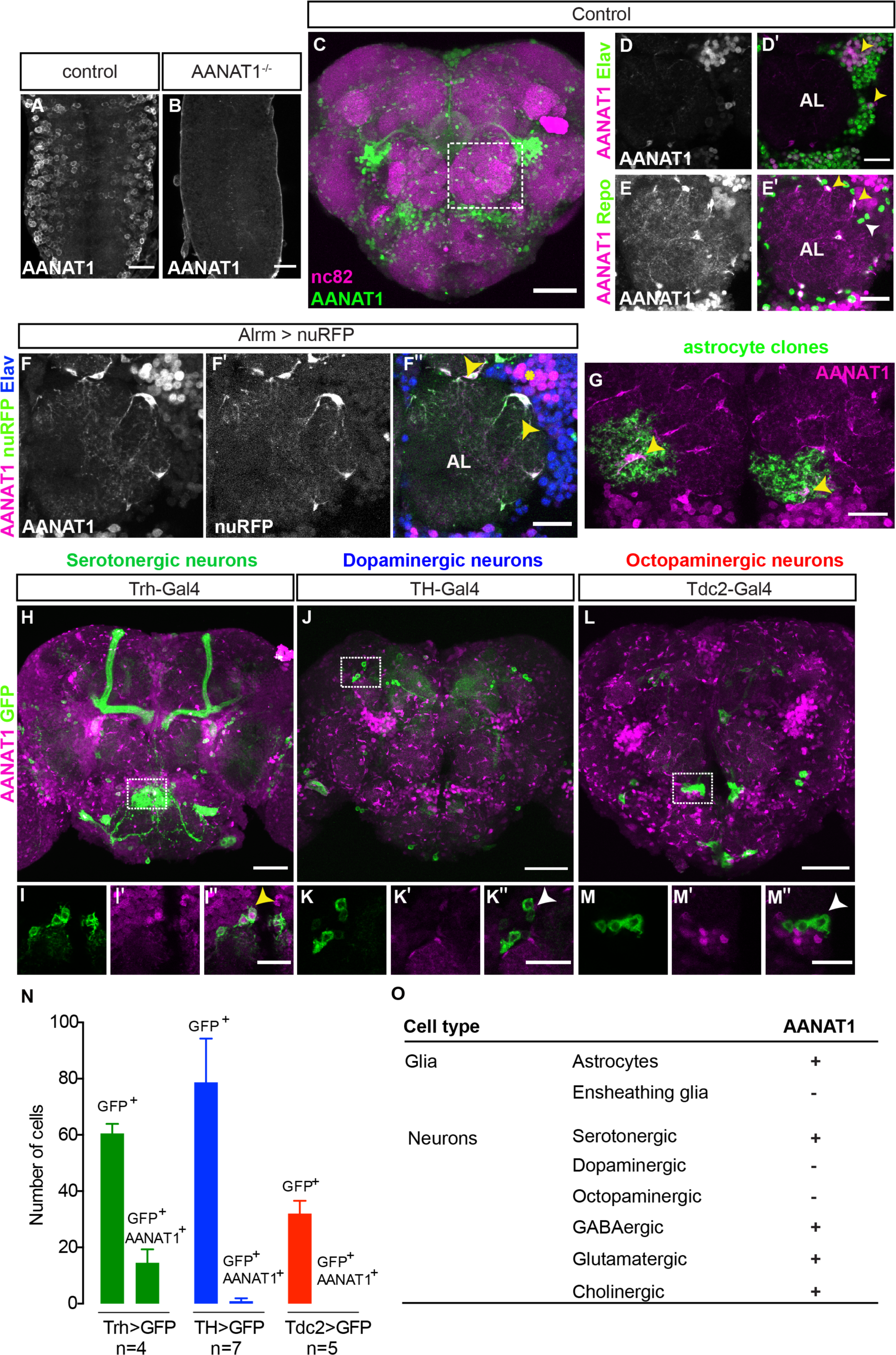
AANAT1 expression in the adult *Drosophila* brain. (A-B) AANAT1 IHC in age-matched embryos (stage 17) from *w^1118^* controls (A) and AANAT1 nulls homozygous Df(BSC)356 (B). (C-M) AANAT1 IHC in the central brain of adults. (C) Z-stack projection showing AANAT1 (green) and neuropil marker nc82 (magenta) in Alrm-Gal4/+ control animals. (D-D’) Single optical slice showing AANAT1 (magenta) and the pan-neuronal marker Elav (green). Yellow arrowheads point to neurons co-expressing both (D’). (E-E’). Single optical slice of AANAT1 (magenta) and the pan-glial marker Repo (green) in control animals where most glia express AANAT1 (E’; yellow arrowheads), but not all (E’; white arrowheads). AL = antennal lobe. (F-F’’) Single slice of AANAT1 (magenta), Elav (blue), and astrocyte marker Alrm-Gal4; UAS-nuRFP (Alrm>nuRFP, green) showing co-expression of AANAT1 and nuRFP in astrocytes (F’’; yellow arrowheads) and with Elav (F’’; yellow asterisk). (G) MCFO-labeled single cell astrocyte clones (green) co-labeled with AANAT1 (magenta). Yellow arrowheads indicate AANAT1-positive astrocyte cell bodies. (H-M’’) Z-stack projections and single slice images of AANAT1 (magenta) and GFP (green) IHC in monoaminergic neurons labeled with type-specific Gal4 drivers. Dotted boxes in H, J and L show regions approximating those selected for imaging at higher power in animals of the same genotypes shown in I, K and M, respectively. AANAT1 is expressed in some serotonergic neurons (I’’; yellow arrowhead), but not in dopaminergic or octopaminergic neurons (K’’, M’’; white arrowheads). (N) Quantification of the mean number of GFP-positive and GFP/AANAT1 double-positive cells in the central brains of animals where Gal4 is used to express GFP in serotonergic (green), dopaminergic (blue) or octopaminergic (red) neurons. Error bars represent standard deviation. (O) Summary of AANAT1 expression in cell types of the adult *Drosophila* central brain. Scale bars in A, B, D-G, I, K, M = 20 µm. Scale bars in C, H, J, L = 50μm.

Labeling of astrocytes was confirmed to be specific for AANAT1 because it was lost upon knockdown of AANAT1 from astrocytes with Alrm-Gal4 or Repo-Gal4 (**Fig. S1**). This also revealed more clearly the several clusters of AANAT1-positive neurons and their axon tracts in the central complex of the brain (**Fig. S1**), which we examined in brain regions associated with sleep regulation (**Fig. S1**). AANAT1 expression was largely absent from the neuropils of the mushroom body and fan-shaped body (FSB), though there were scattered AANAT1-positive astrocytes nearby. AANAT1 expression in the neuropil of the ellipsoid body (EB) came almost exclusively from neurons (**Fig. S1)**. Elsewhere, AANAT1 expression came primarily from the infiltrative processes of astrocytes, as in the neuropil regions of the antennal lobe and subesophageal ganglion. Overall, it appeared that astrocytes contributed far more to AANAT1 labeling of brain neuropil than did neurons.

The monoamines serotonin, dopamine, and octopamine (the insect homolog of norepinephrine) are known to act in the fly brain to regulate the quantity and timing of sleep [2]. Pharmacological, genetic, and thermogenetic approaches have converged to demonstrate that serotonin signaling in the fly brain increases sleep, whereas dopamine or octopamine signaling promote waking [2, 12, 18–22]. Previous studies have suggested AANAT1 expression in dopaminergic neurons [23, 24], but this has not been tested directly. With IHC, we examined AANAT1 co-labeling of serotonergic, dopaminergic and octopaminergic neurons using a mCD8-GFP reporter driven by either Trh-Gal4 [25], TH-Gal4 [26], or Tdc2-Gal4 [27], respectively. AANAT1 was expressed in an average of 14.5 ± 4.8 of 60 ± 3.4 (25%) of serotonergic cells labeled with Trh>mCD8-GFP (**Fig. 1H, I-I’’,N**). AANAT1^+^ serotonergic neurons belonged to a medial subeosophageal ganglion cluster. However, AANAT1 did not co-label cells expressing TH>mCD8-GFP or Tdc2>mCD8-GFP (**Fig. 1J, K-K’’, L, M-M’’, N**), indicating AANAT1 is not expressed in dopaminergic or octopaminergic neurons. These results are corroborated by single-cell RNA sequencing data showing AANAT1 transcripts in astrocytes and serotonergic neurons [28].

To identify the other types of neurons expressing AANAT1, we used a mCD8-GFP reporter driven by either MiMIC-vGlut, Gad1-Gal4, or Cha-Gal4 and found AANAT1 in subpopulations of neurons that release glutamate, gamma-aminobutyric acid (GABA), or acetylcholine, respectively (**Fig.** S**1)**. Monoamines are mainly synthesized in the neurons that release them, and it is generally understood that their re-uptake into these same neurons occurs via specific transport proteins to prevent their accumulation at synapses [29]. Absence of AANAT1 from dopaminergic or octopaminergic neurons showed that cells that produce and release monoamines do not necessarily contribute to their catabolism via AANAT1. However, the presence of AANAT1 in subsets of glutamatergic, GABAergic and cholinergic neurons suggests that, along with astrocytes, these non-monoaminergic neurons could contribute to regulation of monoamine bioavailability in the brain.

The *AANAT1* gene produces two isoforms, the shorter of which (FlyBase AANAT1-PA, 240aa in length), previously known as aaNAT1b, is more predominant [30]. This shorter isoform was observed to be lost in *AANAT1^lo^* mutants [31]. *AANAT1^lo^* is a spontaneous mutation that arose from insertion of a transposable element into the *AANAT1* gene, and extracts of these flies have reduced AANAT1 activity [31, 32]. Using our new AANAT1 antiserum to perform Western blotting of brain extracts, we observed only the short isoform in controls. In *AANAT1^lo^* homozygotes and hemizygotes (*AANAT1^lo^*/In(2LR)Px[4]), AANAT1 protein levels were reduced to 13% and 8% of iso31 controls, respectively (**Fig. 2A,B**). This was confirmed with IHC in the brains of *AANAT1^lo^* flies (**Fig. 2C-E)**, where we noted residual AANAT1 expression in some Elav^+^ neurons, but complete loss of AANAT1 from astrocytes (**Fig. 2F-F’, G-G’**).

**Figure 2.**
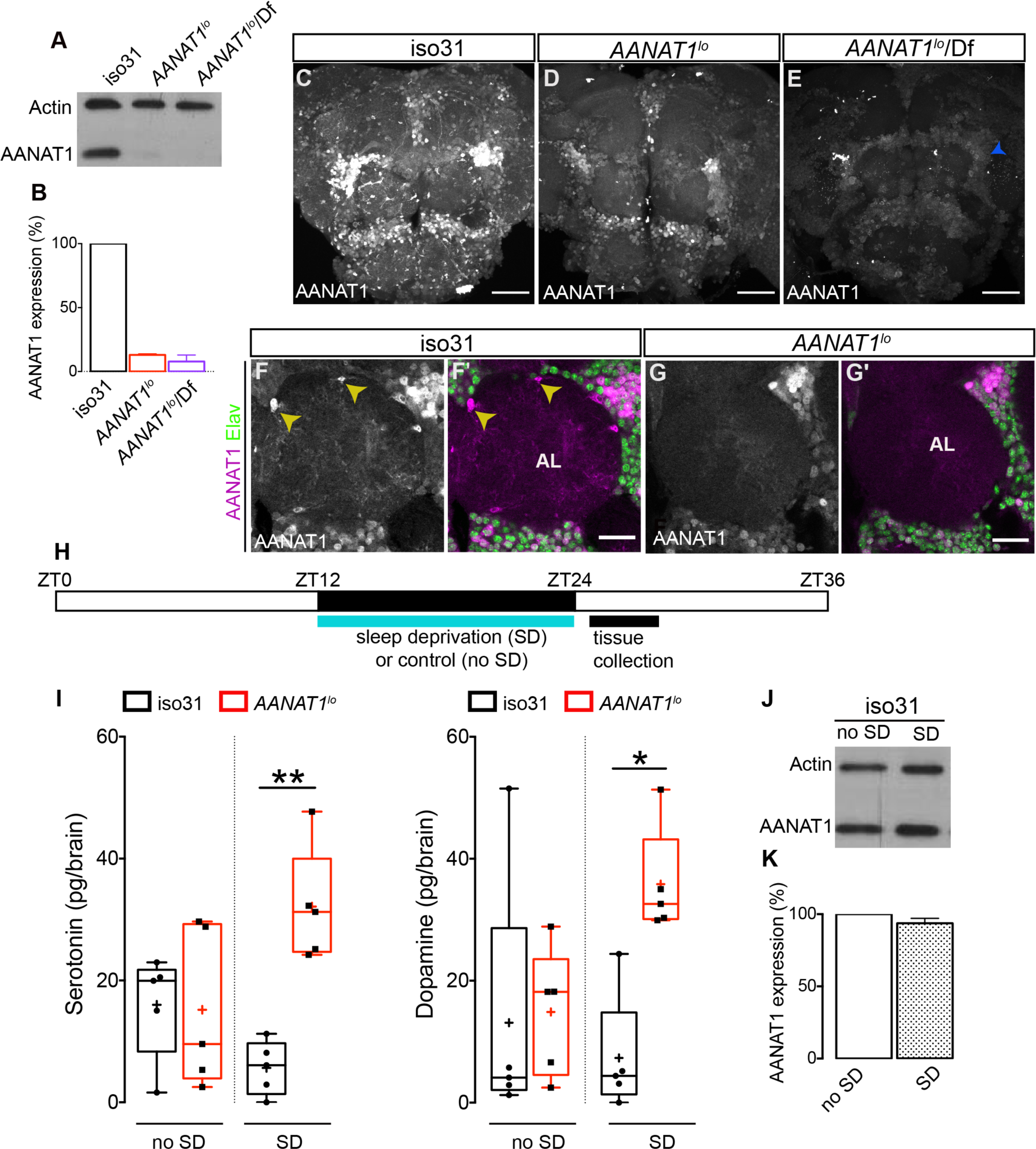
Characterization of *AANAT1^lo^*. (A) Western blot of lysates prepared from dissected brains (ZT15-16) of iso31, *AANAT1^lo^* and *AANAT1^lo^*/Df(In(2LR)Px[4]) adult males. (B) Quantification of AANAT1 expression normalized to that of Actin (mean + standard deviation, n=3 biological replicates). (C-E) Z-stack images showing AANAT1 (grey) in iso31 (C), *AANAT1^lo^* (D) and *AANAT1^lo^*/Df(In(2LR)Px[4]) (E) animals. Blue arrowhead in E represents background signal. Scale bars=50 µm. (F, G) Single optical slices showing AANAT1 (gray or magenta) and Elav (green) in iso31 (F, F’) and *AANAT1^lo^* (G, G’). Yellow arrowhead shows AANAT1^+^ astrocytes. Scale bars = 20 µm. (H) Schematic of experiment for HPLC-MS analysis. (I) HPLC-MS measurement of serotonin (One-way ANOVA with Tukey’s post-hoc test, *p<0.05, **p<0.01,) and dopamine (Kruskal-Wallis test, Dunn’s multiple comparisons, *p<0.05,) in iso31 (black) and *AANAT1^lo^* (red) fly brains under control and sleep deprivation (SD) conditions. Box and whisker plots in this figure show 25-75% interquartile range (box), minimum and maximum (whiskers), median (horizontal line in box), and mean (+). n=5 per genotype. (J) Western blot of lysates prepared from dissected brains (ZT24-25) of iso31 females in control (no SD) and sleep deprivation (SD) conditions. (K) Quantification of AANAT1 (paired t-test, p=0.0831, n=3) expression, normalized to actin levels in iso31 animals under control (no SD) and sleep deprivation (SD) conditions.

*In vitro* studies have shown that serotonin and dopamine are substrates for AANAT1 with similar affinities [17]. Whether the levels of these and/or other monoamines are regulated by AANAT1 *in vivo* remains to be determined. We used HPLC-MS to measure levels of serotonin, dopamine, and octopamine in the brains of *AANAT1^lo^* flies and controls (iso31) (**Fig. 2H**). Under baseline sleep-cycle conditions, where brain tissues were collected in a 3-hour window after lights-ON (ZT0), serotonin and dopamine levels in *AANAT1^lo^* flies were similar to controls (**Fig. 2I**). Octopamine was undetectable in controls, and found at low levels in brains of *AANAT1^lo^* flies (**Fig. S2**). However, if this window was preceded by 12 hours (ZT12-ZT24) of sleep deprivation, *AANAT1^lo^* brains had a robust increase in the levels of serotonin and dopamine compared to controls (**Fig. 2I**), but this had no effect on octopamine levels (**Fig. S2**). Importantly, in control animals the sleep deprivation protocol itself did not appear to affect the levels of measured monoamines, or of AANAT1 itself (**Fig. 2J, K**). Further, we did not observe changes in the levels of another monoamine catabolic enzyme known to be expressed in astrocytes (Ebony) in either *AANAT1^lo^* flies, or in flies subjected to sleep deprivation (**Fig. S2**). We conclude that catabolism of serotonin and dopamine in the brains of flies lacking AANAT1 is severely compromised upon sleep deprivation, leading to inappropriate accumulation of these monoamines.

*AANAT1^lo^* increases homeostatic sleep following deprivation [16], suggesting AANAT1 could be key to how the brain limits the response to sleep deprivation. *AANAT1^lo^* is also interesting because these flies have normal activity, and intact daily patterns of sleep [16], allowing genetic dissection of homeostatic sleep control independent of the regulation of baseline sleep. We wondered whether the increased rebound sleep seen in *AANAT1^lo^* animals could be explained by loss of AANAT1 function from neurons or astrocytes. To test this, we selectively knocked down AANAT1 in distinct cell types with RNAi, and measured both baseline and homeostatic sleep with the *Drosophila* Activity Monitoring System (DAMS). To evaluate the contribution of neuronal AANAT1 in sleep, we tested nSyb-Gal4>UAS-AANAT1-RNAi flies and found that they displayed normal patterns of baseline sleep (**Fig. 3A-H**), as reported for the *AANAT1^lo^* allele [16]. However, knockdown of AANAT1 in neurons increased the amount of nighttime sleep compared to controls (**Fig. 3B, F**). In addition, AANAT1 knockdown in neurons led to sleep bouts during the night that were increased in duration and decreased in number, suggesting improved sleep consolidation at night (**Fig. 3C-D;G-H**). This was observed for two independent RNAi lines that target the AANAT1 transcript at distinct sites, though the effect was stronger for one (AANAT-RNAi 2, JF02142) than the other (AANAT-RNAi 1, HMS01617). We then assessed whether AANAT1 knockdown in neurons (nSyb>AANAT1-RNAi) would impact sleep rebound after sleep deprivation, as was observed in *AANAT1^lo^* flies. For this, flies were subjected to overnight mechanical sleep deprivation and we found, somewhat surprisingly, that these flies do not display an enhanced rebound sleep the next day (**Fig. 3I-L**). Flies lacking AANAT1 in neurons were not totally deprived of sleep by the mechanical stimulation, but as this was also the case in one of the controls, it is not indicative of a phenotype of AANAT1 loss.

**Figure 3.**
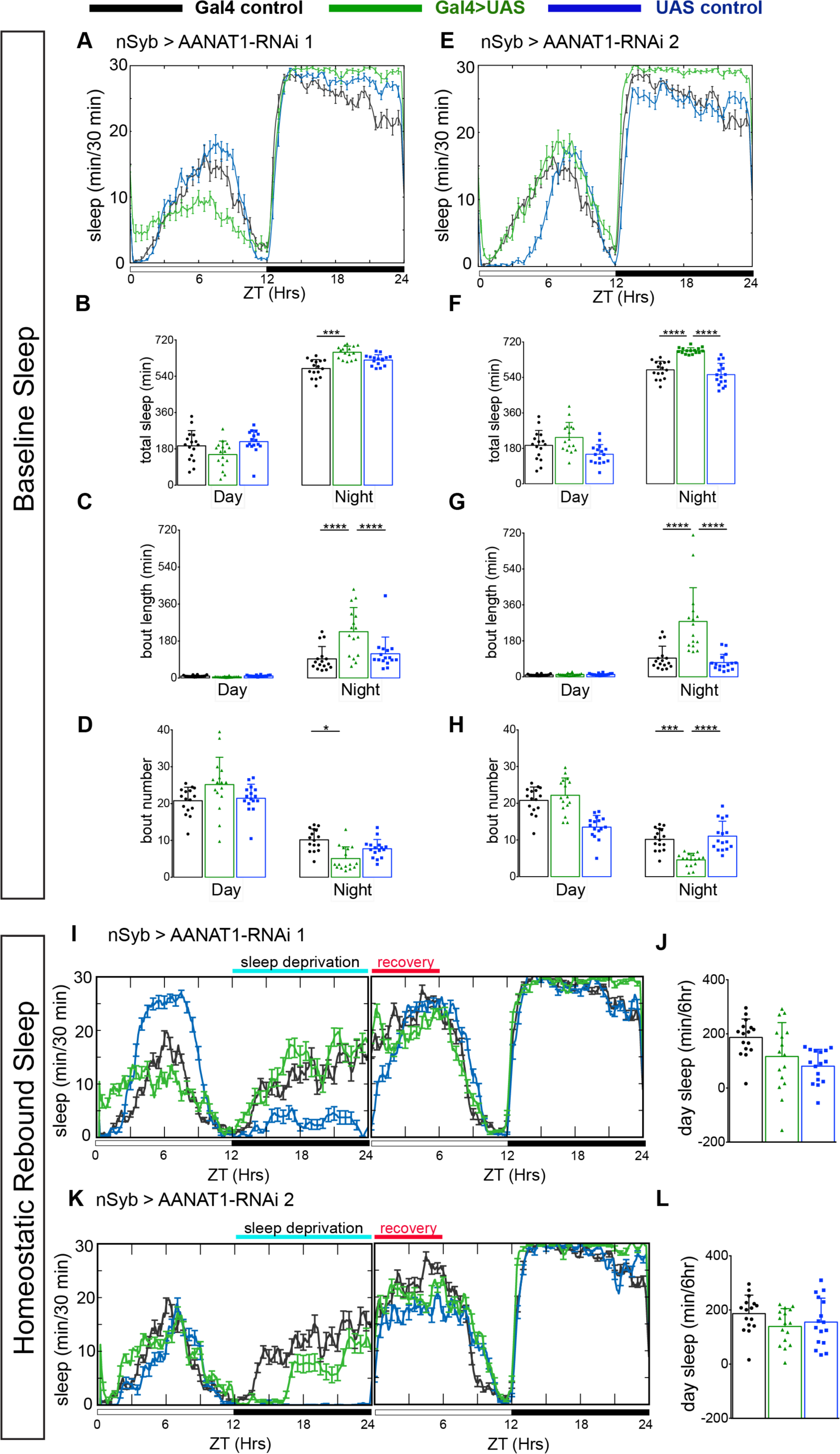
AANAT1 knockdown in neurons. (A-D) Baseline sleep upon knockdown with UAS-HMS01617 (RNAi 1). 24-hr sleep profile showing light/dark conditions on X-axis (A), and quantification during day (ZT 0-12) versus night (ZT 12-24) of total sleep duration (B), sleep bout length (C) and bout number (D) for the nSyb-Gal4 control (black), the UAS-HMS01617 control (blue), and knockdown animals (nSyb>HMS01617, green). (n=16 per genotype, bar graphs show mean + standard deviation, one-way ANOVA with Tukey’s post-hoc test, *p<0.05, ***p<0.001, ****p<0.0001). (E-H) Baseline sleep upon knockdown with UAS-JF02142 (RNAi 2). (E) 24-hr sleep profile showing light/dark conditions on X-axis. Quantification of total sleep duration (F), sleep bout length (G) and bout number (H) for the nSyb-Gal4 control (black), the UAS-JF02142 control (blue), and knockdown animals (nSyb>JF02142, green). The plotted nSyb-Gal4 control data is the same as in A-D, as the experiments were done simultaneously. (n=16 per genotype, one-way ANOVA with Tukey’s post-hoc test, ***p<0.001, ****p<0.0001). (I, J) Rebound sleep upon knockdown with UAS-HMS01617 (RNAi 1). (I) 24-hr sleep profile of baseline and recovery days, and (J) the duration of sleep during ZT0-6 recovery period. nSyb-Gal4 control (black), the UAS-HMS01617 control (blue), and knockdown animals (nSyb>HMS01617, green). (n=16 per genotype, two-way ANOVA with Tukey’s post-hoc test). (K, L) Rebound sleep upon knockdown with UAS-JF02142 (RNAi 2). (K) 24-hr sleep profile of baseline and recovery day, (L) duration of sleep during ZT0-6 recovery period (L). nSyb-Gal4 control (black), the UAS-JF02142 control (blue), and knockdown animals (nSyb>HMS01617, green). (n=16 per genotype, two-way ANOVA with Tukey’s post-hoc test).

Next, we used Alrm-Gal4 to selectively deplete AANAT1 expression from astrocytes with RNAi (Alrm>AANAT1-RNAi). This had no effect on the numbers of astrocytes present in the brain (**Fig. 4A**), and flies showed normal baseline patterns and amounts of daytime and nighttime sleep compared to controls carrying either GAL4 or UAS transgenes alone (**Fig. 4B-E**), consistent with reported normal circadian sleep-cycle and motor activity of *AANAT1^lo^* [16]. However, upon overnight mechanical sleep deprivation, these flies had increased rebound sleep the next day (**Fig. 4F-I)**, mimicking *AANAT1^lo^* flies. This effect was partially rescued by simultaneous expression of AANAT1-myc in animals where the RNAi targeted the non-coding 3’ UTR of endogenous AANAT1 mRNA (AANAT-RNAi 1 (HMS01617), **Fig. 4J**). Incomplete rescue might be explained by our observation that AANAT1-myc was not detectable in many brain astrocytes in these animals (**Fig. 4K**). Taken together, our results demonstrate that AANAT1 acts in astrocytes, but not in neurons, to restrict daytime rebound sleep in response to overnight sleep deprivation. Along with our finding that AANAT1 limits accumulation of serotonin and dopamine upon sleep deprivation, these data suggest that *Drosophila* astrocytes are specifically engaged to handle particular demands on monoamine metabolism imposed by sleep deprivation overnight.

**Figure 4.**
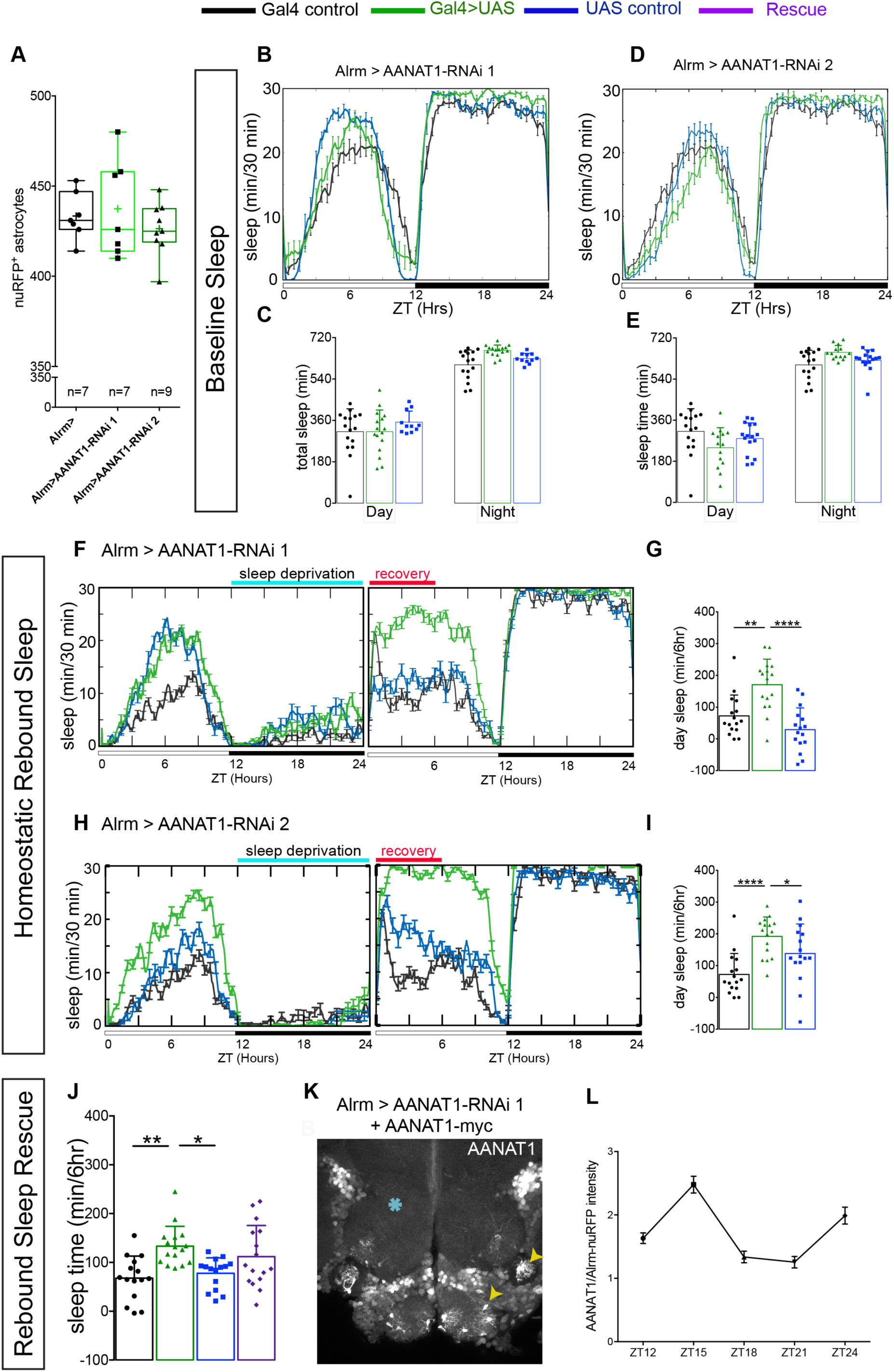
AANAT1 knockdown in astrocytes. (A) Compared with Alrm-Gal4 controls (Alrm>), the number of nuRFP labeled astrocytes in the central brain is unaffected upon RNAi knockdown of AANAT1 with HMS01617 (AANAT1-RNAi 1) or JF02142 (AANAT1-RNAi 2). Box and whisker plot as in Figure 2F. One-way ANOVA with Tukey’s post-hoc test, n=7-9 per genotype. (B, C) Baseline sleep upon knockdown with UAS-HMS01617 (RNAi 1). 24-hr sleep profile (B), and total sleep duration (C) for the Alrm-Gal4 control (black; n=16), the UAS-HMS01617 control (blue; n=11), and knockdown animals (Alrm>HMS01617, green; n=16). (One-way ANOVA with Tukey’s post-hoc test). (D, E) Baseline sleep upon knockdown with UAS-JF02142 (RNAi 2). 24-hr sleep profile (D), and total sleep duration (E) for the Alrm-Gal4 control (black; n=16), the UAS-JF02142 control (blue; n=16), and knockdown animals (Alrm>JF02142, green; n=14). The plotted Alrm-Gal4 control data is the same as in B and C, as the experiments were done simultaneously. (One-way ANOVA with Tukey’s post-hoc test). (F, G) Rebound sleep upon knockdown with UAS-HMS01617 (RNAi 1). 24-hr sleep profile of baseline day and recovery day (F), and the duration of sleep during ZT0-6 recovery period (G) for the Alrm-Gal4 control (black), the UAS-HMS01617 control (blue), and knockdown animals (Alrm>HMS01617, green). (n=16 per genotype, two-way ANOVA with Tukey’s post-hoc test, **p<0.01, ****p<0.0001). (H, I) Rebound sleep upon knockdown with UAS-JF02142 (RNAi 2). 24-hr sleep profile of baseline day and recovery day (H), and the duration of sleep during ZT0-6 recovery period (I) for the Alrm-Gal4 control (black), the UAS-JF02142 control (blue), and knockdown animals (Alrm>HMS01617, green). (n=16 per genotype, error bars are mean + standard deviation, Two-way ANOVA with Tukey’s post-hoc test, *p<0.05, ****p<0.0001). (J) Rebound sleep rescue with UAS-AANAT1-myc in the Alrm-Gal4 control (black), the UAS-HMS01617 control (blue), knockdown animals (Alrm>HMS01617, green) and rescue (Alrm>HMS01617 + UAS-AANAT1-myc, purple). (n=16 per genotype, error bars are mean + standard deviation, One-way ANOVA with Tukey’s post-hoc test, *p<0.05, **p<0.01). (K) Z-stack projections of Alrm>HMS01617 + AANAT1-myc, where AANAT1 (grey) expression is incompletely restored to astrocytes (yellow arrowheads). Blue asterisks indicate AANAT1-negative neuropil region, validating the efficacy of RNAi. (L) AANAT1 levels in astrocyte cell bodies normalized to nuRFP at ZT12,15,18,21 and 24 time-points. (n=3 per time-point, 10 cells per sample, mean+SEM).

With IHC we examined AANAT1 under baseline sleep-cycle conditions at 3 hr intervals during the dark period ZT12-ZT24. Interestingly, we found that AANAT1 expression in astrocytes peaked at ZT15, declined markedly to undetectable levels by ZT21 (**Fig. 4L**), and was re-established at lights-ON (ZT24). From this we infer that the loss of AANAT1 might have more profound effects during ZT12-15 of sleep deprivation, or during daytime afterward, though further temporal refinement of astrocytic AANAT1 function is required. AANAT1 is expressed in astrocytes that reside throughout the brain, and so it remains unclear whether it modulates sleep homeostasis by acting within a particular region of the brain, or more broadly. Interestingly, the neuropils of key sleep centers (EB and FSB; **Fig. S1**) had no AANAT1 staining from infiltrative astrocytes, raising the likelihood it acts elsewhere. Finally, it remains to be established whether the effect of AANAT1 on sleep homeostasis is due to serotonin, dopamine, or both. In *Drosophila*, serotonergic signaling in the brain promotes sleep, while dopaminergic signaling promotes waking. Levels of both serotonin and dopamine are upregulated in *AANAT1^lo^* mutants upon sleep deprivation, where increased sleep prevails. It stands to reason that AANAT1 could act in astrocytes to limit the deprivation-dependent accumulation of sleep-promoting serotonin. It is also possible that dopamine accumulation plays a role, since thermogenetic activation of wake-promoting dopaminergic neurons at night promotes compensatory sleep the next day. This suggests these particular neurons are upstream of circuits that produce homeostatic responses to extended wakefulness [13, 33], and astrocytic AANAT1 could somehow restrict dopaminergic signaling from these neurons overnight.

Our findings illustrate a newly-discovered role for astrocytes in the control of monoamine bioavailability and homeostatic sleep drive. *Drosophila* astrocytes also express the enzyme Ebony, which couples dopamine to N-β-alanine [34], and a receptor for octopamine and tyramine [35], reinforcing how they are well-equipped to metabolize monoamines, and to monitor and respond to monoaminergic neuronal activity. Neither gene expression studies nor RNA sequencing databases provide evidence for monoamine-synthesizing enzymes in *Drosophila* astrocytes, so it appears likely that monoamines inactivated by AANAT1 in astrocytes are brought into these cells by an unidentified transporter. Astrocytes are particularly well-suited for regulating sleep in this way because they have ramified processes that infiltrate neuropil regions to lie in close proximity to synapses.

AANAT1 also functions in neurons to regulate baseline sleep, which highlights the idea that cellular context has profound impact on AANAT1 function in sleep regulation. In light of this, we note that loss of the related enzyme *aanat2* in zebrafish larvae decreases baseline sleep [36], which could be attributed to a loss of melatonin since the AANAT1 product N-acetylserotonin is an intermediate in the synthesis of melatonin in vertebrates [31]. Clearly, the appropriate balance and cellular context of AANAT activity is critical for the regulation of sleep, and we show here in *Drosophila* that astrocytes are an important contributor to this balance. Interestingly, astrocytes in rodents express the monoamine transporters and receptors for dopamine and serotonin [37–42], raising the possibility that astrocytes in mammals might also participate in mechanisms of sleep regulation involving monoaminergic neural signaling.

## Methods

### Fly stocks

*Drosophila melanogaster* stocks were obtained from the Bloomington *Drosophila* Stock Center (BSC): Trh-Gal4 (BSC-52249), TH-Gal4 (BSC-8848), Tdc2-Gal4 (BSC-9313), Ddc1-Gal4 (BSC-7010), UAS-mCD8-GFP (BSC-32186), UAS-RFP.nls (BSC-30558), Mi{MIC} VGlut^MI04979^ (BSC-38078), Gad1-Gal4 (BSC-51630), Cha-Gal4 (BSC-6793), R56F03-Gal4 (BSC-39157), *AANAT1^lo^* (BSC-3193), Df(2R)BSC356 (BSC-24380), deficiency In(2LR)Px4 (BSC-1473), AANAT1 RNAi lines UAS-HMS01617 (BSC-36726), UAS-JF02142 (BSC-26243) and multi-color Flp-out stock hs-FlpG5.Pest; 10xUAS(FRT-stop)myr::smGdP-HA, 10xUAS(FRT-stop)myr::smGdP-V5-THS-10xUAS(FRT-stop)myr::smGdP-FLAG (BSC-64085). Alrm-Gal4 was provided by Dr. Marc Freeman, and nSyb-Gal4 by Dr. Stefan Thor.

For RNA interference (RNAi) knockdown of gene expression, control animals carried only a Gal4 driver, while experimental groups also carried a single copy of the transgene to elicit RNAi. The chromosome carrying Alrm-Gal4 also bore a transgene encoding the nuclear reporter UAS-nuRFP. To mitigate the effects of genetic background for sleep experiments, control Gal4 and UAS flies were crossed to the iso31 stock.

The morphology of single astrocytes was determined by the Multi-Color Flp-OUT (MCFO) technique [43], where three differently tagged reporters under UAS control (HA, FLAG and V5) were silenced by FRT-flanked transcriptional terminators. Heat shock-induced FLPase expression removed terminators randomly in individual cells, driven by astrocyte-specific Alrm-GAL4. This created a mosaic of differently-colored astrocytes. For this experiment, 3-5 day old flies raised at 18°C were heat-shocked at 37°C for 5-8 minutes and dissected 2–3 days later.

To create UAS-AANAT1-myc, AANAT1 coding sequence from cDNA clone GH12636 (*Drosophila* Genomic Research Centre) was PCR-amplified and cloned in-frame into a modified pJFRC-MUH [44] carrying a C-terminal myc-tag. Transgenic flies with site-specific insertions at *VK0005* site on chromosome 3 were generated using standard microinjection (BestGene, Inc.).

### Generation of AANAT1 antibody

A KLH-coupled peptide RRPSPDDVPEKAADSC (amino acids (aa) 94-109 of isoform AANAT1-PA (FlyBase), or 129-144 of isoform AANAT1-PB) was synthesized and injected into rabbits according to guidelines of the Canadian Council for Animal Care (MEDIMABS, Montreal, QC).

### Immunohistochemistry and imaging

Adult fly brains were dissected in cold phosphate-buffered saline (pH 7.4) and fixed in 4% paraformaldehyde for 30 minutes (min). After three washes of 15 min each with PBS containing 0.3% Triton-X-100 (PBTx-0.3%), the tissues were blocked in 5% normal goat serum (Jackson Laboratories) in PBTx-0.5% for 45 min. Tissues were incubated in primary antibodies: rabbit anti-AANAT1 (1:2000; this study), rat anti-Elav (1:100; Developmental Studies Hybridoma Bank (DSHB), mouse anti-nc82 (1:50; DSHB), mouse anti-GFP (1:200; Clontech #632381) overnight at 4⁰C. After three washes (15 min each, PBTx-0.3%), tissues were incubated with secondary antibodies overnight at 4⁰C: goat anti-mouse (Rhodamine Red-X, Jackson ImmunoResearch #115-295-146), goat anti-rabbit (Alexa Fluor 488, Thermo Fisher Scientific, #A11008), goat anti-mouse (Alexa Fluor 488, Thermo Fisher Scientific), goat anti-rat (Alexa Fluor 568, Thermo Fisher Scientific, #A11077), goat anti-rabbit (Alexa Fluor 647, Thermo Fisher Scientific, #A21245). Tissues were again washed (3 × 15min, PBTx-0.3%), followed by a final wash in PBS. Tissues were mounted in SlowFade™ Diamond Antifade Mountant (Thermo Fisher Scientific, #S36964). Fluorescence images were acquired with an Olympus BX-63 Fluoview FV1000 confocal laser-scanning microscope and processed using Fiji.

For MCFO labeling, brains were dissected in ice-cold PBS, fixed with 4% paraformaldehyde/PBS for 1 h at room temperature followed by three successive washes in 0.5% PBTx for 20 min each. Simultaneous incubation (48 h at 4 °C) with rat anti-FLAG (1:100; Novus Biologicals NBP1-06712,A-4) and rabbit anti-AANAT1 was followed by another 48 h at 4 °C with goat anti-rabbit (1:1000; Alexa Fluor 488, Thermo Fisher Scientific, #A11008), goat anti-rat (1:1000; Alexa Fluor 568, #A11077) and V5-tag:AlexaFluor-647 (1:200; Bio-Rad MCA1360A647).

To quantify cells immuno-labelled for GFP and AANAT1, cells were manually counted from image stacks of the central brain (excluding optic lobes).

### Western blotting

Lysates for western blots were prepared from dissected adult brains in 50μl Laemmli buffer as reported in [45]. 10 brains were used per lysate and incubated at 90⁰C for 5 min. 15μl of each sample was loaded per well, run on 15% SDS-PAGE gels, blotted to nitrocellulose membrane, and probed with Rabbit anti-AANAT1 (1:2500) or anti-Ebony (1:3000; Sean Carroll, University of Wisconsin-Madison), and mouse anti-actin (1:3000; Sigma #A4700). HRP-conjugated secondary antibodies anti-rabbit (1:3000; Bio-Rad) and anti-mouse (1:3000; Promega #W4021) were used for detection with chemiluminescence (HyGLO Chemiluminescent HRP Antibody Detection Reagent, Denville Scientific). Mean signal intensity for AANAT1 or Ebony was quantified using Fiji and normalized to actin. We used three separate lysates for each genotype to analyze western blots. For sleep experiments, female brains were used for lysate preparation.

### High Performance Liquid Chromatography – Mass Spectrometry (HPLC-MS)

To prepare samples for HPLC-MS, the brains of twenty female flies (1-2 weeks old) for each genotype were dissected into ice-cold PBS between ZT0.5-3.5. We dissected brain tissue to avoid cuticle contamination because serotonin and dopamine are intermediates in the sclerotization of *Drosophila* cuticle. Dissected brains were centrifuged, the PBS was removed, and samples were quickly homogenized with a motorized pestle into an aqueous solution of formic acid (0.1%). After centrifugation, the supernatant was collected and stored at −80°C. Preliminary analytical conditions were developed using reference standards in a solution containing either serotonin, dopamine, or octopamine. With LC-MS/MS (Thermo-Scientific Quantiva Triple Quadrupole Mass Spectrometer (QQQ)), the absolute values for each analyte were measured in picograms (pg) per brain, through the addition of deuterated reference standards to sample extracts.

### Monitoring and measurement of sleep in *Drosophila*

Prior to experimentation, flies were kept on standard food in constant conditions (a 12-hour light/dark cycle, and 25°C). At least 5 days after eclosion, mated adult females were loaded into glass tubes with 5% sucrose/2% agar food for behavioral recordings. The *Drosophila* Activity Monitoring (DAM) system (Trikinetics, Waltham, MA) was used to quantify infrared beam breaks representing locomotor activity. Files were processed with PySolo [46] in 1-minute bins, with sleep defined as 5 consecutive minutes without activity, as done previously [47]. In sleep deprivation experiments, flies were placed in DAM monitors on a vortexer that was mechanically shaken a random 2 of every 20 seconds over the course of the 12 hours of the dark period (ZT12-24). Rebound sleep was determined, per fly, as the difference between sleep amount in the period following deprivation and sleep amount in the same time period on the preceding baseline day in unperturbed conditions.

### Time-course of AANAT1 expression in astrocytes

AANAT1 levels in astrocytes were quantified at 3 hr intervals between ZT12-24 with IHC, where AANAT1 fluorescence intensity in astrocyte cell bodies was measured and normalized to nuRFP intensity from 2 copies of a UAS-nuRFP transgene reporter driven by Alrm-Gal4. At each time point, 10 astrocytes in the antennal lobe region were measured from each of three brains.

## Acknowledgements

We thank Dr. Marc Freeman (Oregon Health & Science University) and Dr. Sean Carroll (University of Wisconsin-Madison) for reagents, to members of the van Meyel lab for helpful critique, and especially to Dr. Emilie Peco, Dr. Tiago Ferreira, Dr. Yimiao Ou, and Renu Heir for advice and assistance. LC-MS analysis was performed at the Proteomics Platform of the Research Institute of McGill University Health Centre.

## Author contributions

SD, GA, AS and DvM contributed to the experimental design, data analysis, and writing of the manuscript. Experiments were performed by SD, GA, DC, and SL.

## Declaration of Interests

The authors declare no competing interests.

## Supplemental Information

**Figure S1, Related to Figure 1. AANAT1 expression in the adult *Drosophila* brain.**

(A) Single optical slice showing AANAT1 (magenta), Elav (blue) and Alrm>nuRFP (green), with AANAT1-negative astrocytes (white arrowheads) in the optic lobe.

(B) Single optical slice showing AANAT1 (magenta) and R56F03>mCD8-GFP (green) showing absence of AANAT1 in ensheathing glia (white arrowheads).

(C-F) AANAT1 expression upon AANAT1 knockdown with UAS-HMS01617 RNAi driven in all glia (C; Repo-Gal4;UAS-Dcr-2), all neurons (D; nSyb-Gal4), astrocytes (E; Alrm-Gal4,UAS-nuRFP), and both in neurons and astrocytes (F; nSyb-Gal4;Alrm-Gal4). Cyan arrowheads in C and E depict axonal bundles. Blue arrowhead in F shows background signal.

(G, H) Single slices of IHC for AANAT1 (magenta) and Elav (green) showing loss of AANAT1 expression (white arrowheads) upon knockdown in neurons using nSyb-Gal4 to drive either UAS-HMS01617 (G; RNAi 1) or UAS-JF02142 (H; RNAi 2).

(I, J) Single slices of IHC for AANAT1 (magenta) and Alrm>nuRFP (green) showing loss of AANAT1 expression (white arrowheads) upon knockdown in astrocytes using Alrm-Gal4 to drive RNAi 1 (I) or RNAi 2 (J). Yellow arrowheads indicate AANAT1 expression in neurons.

(K-P) Z-stack projections showing AANAT1 (grey) expression in ellipsoid body (K-M) and fan-shaped body (N-P) neuropil regions in UAS-JF02142/+ controls (K,N) and upon knockdown of AANAT1 with AANAT1-RNAi2 driven in astrocytes using Alrm-Gal4 (L,O) or in neurons using Elav-Gal4 (M,P).

(Q-V’’) Z-stack projections (Q,S,U) and single-slice images (R-R’’;T-T’’;V-V’’) of AANAT1 (magenta) and GFP (green) IHC in non-monoaminergic neurons labeled with type-specific drivers. Dotted boxes in Q, S and U show regions approximating those selected for imaging at higher magnification in animals of the same genotypes shown in R-R’’ T-T’’ and V-V’’, respectively. AANAT1 is expressed in subsets of glutamatergic, GABAergic and cholinergic neurons (R’’, T’’, V’’; yellow arrowheads).

Scale bars in A, C-F, Q, S, U = 50 µm. Scale bars in B, G-J, K-P, R, T, V = 20μm.

**Figure S2, Related to Figure 2. Characterization of *AANAT1^lo^*.**

(A) Quantification of Ebony expression normalized to that of Actin (mean + standard deviation, n=3 biological replicates, One-way ANOVA with Tuckey’s post-hoc test).

(B) HPLC-MS measurement of octopamine (Mann-Whitney test, p<0.533) in *AANAT1^lo^* animals under control (n=2) and sleep deprivation (SD; n=5) conditions. Box and whisker plot as in Figure 2F.

(C) Quantification of Ebony (paired t-test, p=0.7036, n=3) expression normalized to actin levels in iso31 animals under control and sleep deprivation (SD) conditions.

**Figure S4, Related to Figure 4. AANAT1 knockdown in astrocytes.**

(A-A’’) Single optical slice showing AANAT1 (magenta) overexpression in ensheathing glia using R56F03>mCD8-GFP (green).

