## Supplementary figures and images for "AANAT1 functions in astrocytes to regulate sleep homeostasis"

### Supplementary Figures

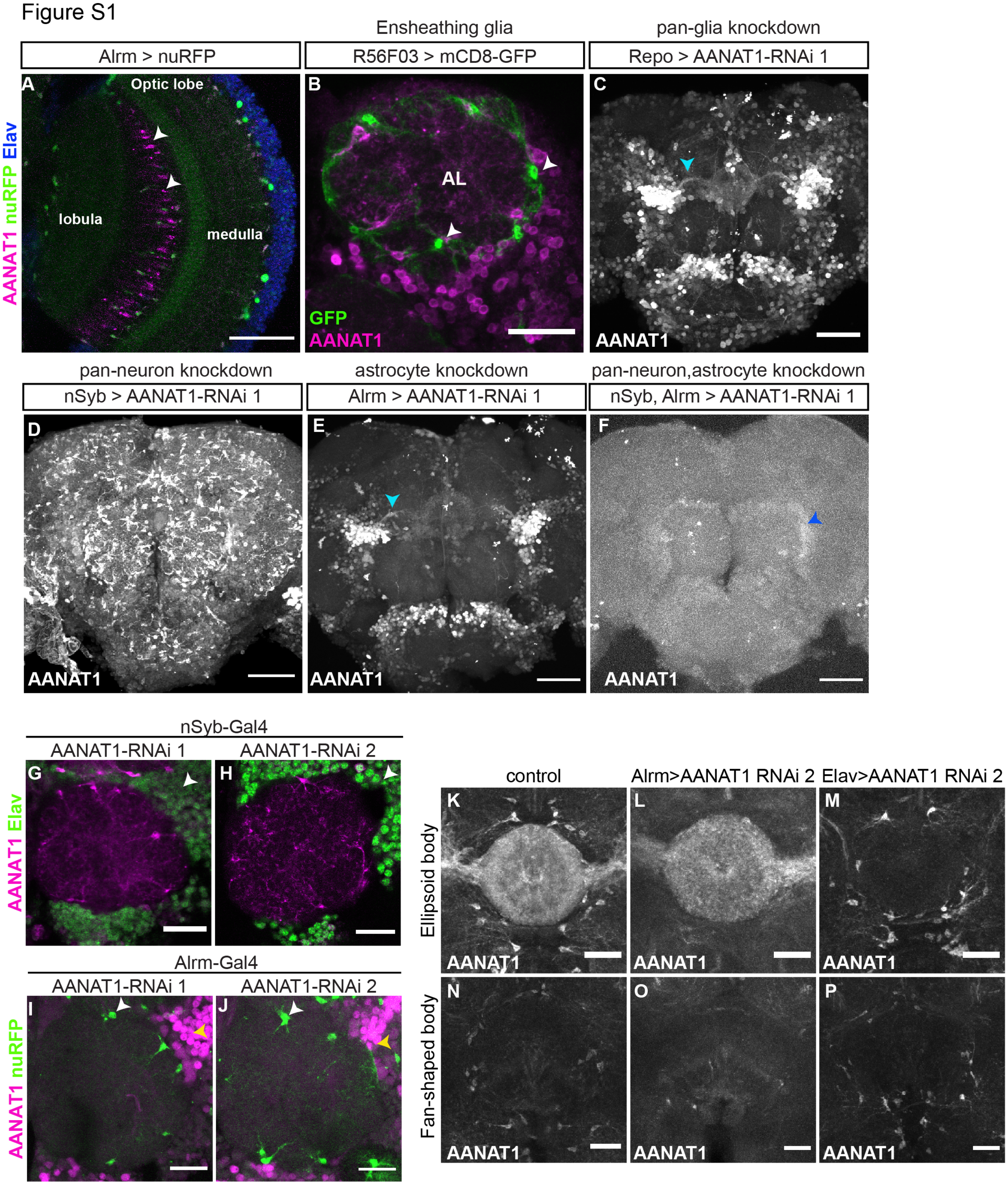

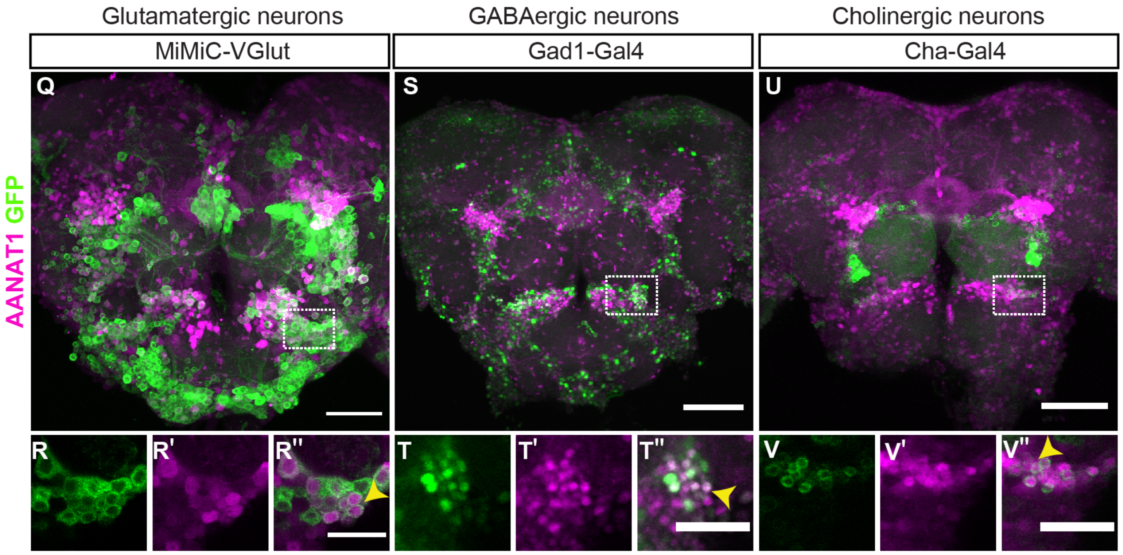

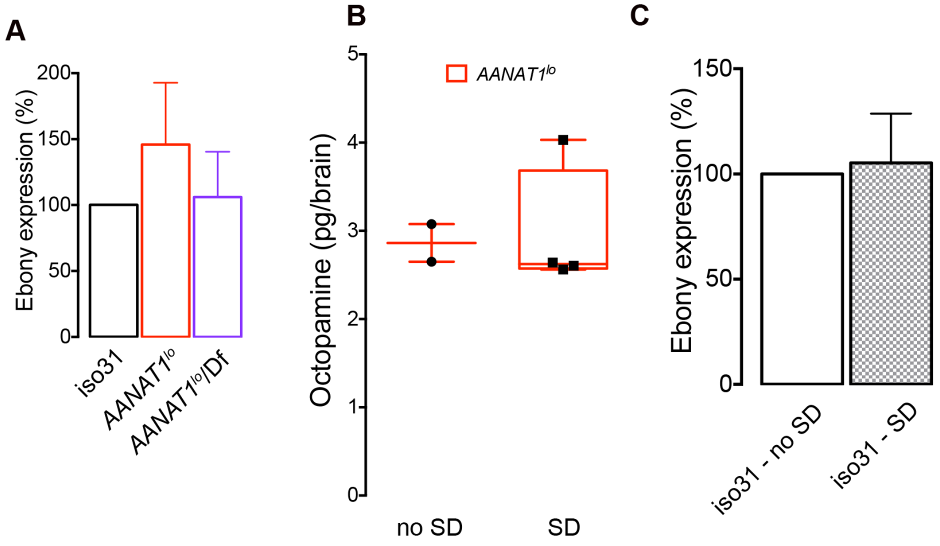

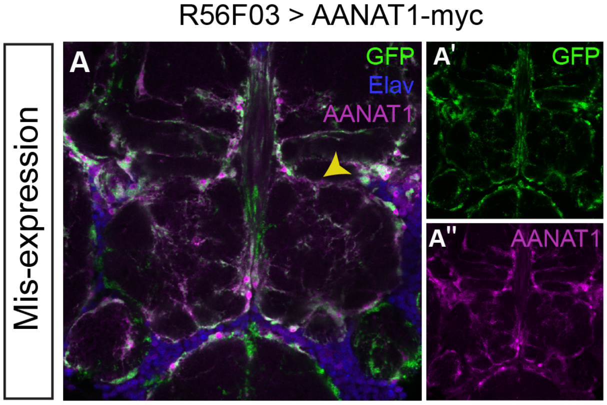
